# C17orf53 defines a novel pathway involved in inter-strand crosslink repair

**DOI:** 10.1101/758722

**Authors:** Chao Wang, Zhen Chen, Dan Su, Mengfan Tang, Litong Nie, Huimin Zhang, Xu Feng, Rui Wang, Xi Shen, Mrinal Srivastava, Megan E. McLaughlin, Glen Traver Hart, Lei Li, Junjie Chen

## Abstract

ATR kinase is a master regulator of genome maintenance and participates in DNA replication and various DNA repair pathways. In a genome-wide screening for ATR-dependent fitness genes, we identified a previously uncharacterized gene, C17orf53, whose loss led to hypersensitivity to ATR inhibition. C17orf53 is conserved in vertebrate and is required for efficient cell proliferation. Loss of C17orf53 slowed down DNA replication and led to pronounced ICL repair defect. Further genetic analyses revealed that C17orf53 functions downstream in ICL repair pathway, probably by affecting the loading of repair factors such as RAD54. In addition, we showed that C17orf53 is a ssDNA- and RPA-binding protein, both of which are important for its functions in the cell. Taken together, C17orf53 is a novel component involved in ICL repair pathway.

## Introduction

The ataxia telangiectasia mutated and RAD3-related (ATR) protein kinase is a master operator in genome maintenance through coordinating DNA replication, DNA repair, and cell cycle transition (de Klein et al., 2000; Hekmat-Nejad et al., 2000; Saldivar et al., 2017). Single-stranded DNA (ssDNA) produced at sites of stressed or stalled replication forks is bound by replication protein A (RPA) complex, which activates ATR with the help of ATR interacting protein (ATRIP) (Bomgarden et al., 2004; Cortez et al., 2001; Zou and Elledge, 2003). The activated ATR further initiate a series of cellular processes including replication stress response and DNA repair. Among these, a critical role of ATR is the control of cell cycle checkpoint. ATR activates its downstream CHK1 kinase, which further suppresses cyclin-dependent kinases (CDKs) to prevent cells with DNA lesions to entry mitosis (Abraham, 2001; Liu et al., 2000; Sorensen and Syljuasen, 2012). A second key function of ATR is to prevent replication catastrophe through stabilizing stressed replication fork (Cha and Kleckner, 2002; Costanzo et al., 2003; Hekmat-Nejad et al., 2000; Luciani et al., 2004). Moreover, ATR activity is important for various DNA damage repair pathways, such as homologous combination repair (HR) and interstrand crosslink (ICL) repair (Buisson et al., 2017; Collis et al., 2008; Fasullo and Sun, 2008; Wang et al., 2004; Zhu and Dutta, 2006).

The multiple functions of ATR in coordinating the responses to replication stress and DNA damage make it a promising target for cancer therapy (Rundle et al., 2017; Williamson et al., 2016). Tumors often harbor high replication stresses due to escalated proliferation, which may increase their need for ATR activity. Similarly, due to the roles of ATR in various DNA damage repair pathways, combination with ATR inhibitor may enhance the efficacy of chemotherapy and radiation therapy that directly or indirectly induce DNA lesions.

DNA interstrand crosslink (ICL) is considered as one of the most toxic DNA lesions because DNA double helix is covalently linked and DNA replication and transcription are absolutely blocked (Huang et al., 1995; Knipscheer et al., 2009). Repair of ICL is a complex process, which requires the coordination of several distinct DNA repair pathways, including Fanconi anemia (FA) pathway, Homologous Recombination (HR) and translesion synthesis (TLS) (D’Andrea, 2001; Huang et al., 2014; Kruyt et al., 1996; Rosselli et al., 2003; Shen et al., 2009; Williams et al., 2013). ATR plays an important role in ICL repair. ATR is activated by ICL-induced stalled replication forks and then in turn phosphorylates numerous downstream effectors, including FA proteins (Shigechi et al., 2012; Tomida et al., 2013). FANCD2 and FANCI proteins are the core components of FA pathway, which acts in part by initiating ICL repair (Joo et al., 2011). Monoubiqitination of FANCI-D2 complex promotes “unhooking” of ICL, which further generates DNA strand breaks at stalled replication forks. HR machinery acts at stalled replication forks to complete DNA repair. HR may also act with TLS to bypass remaining DNA lesions and start DNA replication (Haynes et al., 2015; Hinz, 2010; Roy and Scharer, 2016; Sasaki et al., 2004; Shukla et al., 2013). Since many of these events are regulated by ATR kinase, ATR inhibition sensitizes cells to a variety of DNA damage-inducing agents including cisplatin and mitomycin C (MMC) that induce ICLs.

In order to understanding various cellular processes that require ATR activity in humans, several large-scale genetic screens have been conducted in human cells. Previous studies with RNA interference identified many genes when lost confer heightened sensitivity to ATR inhibition (Kwok et al., 2016; Mohni et al., 2014; Williamson et al., 2016). The advance of CRISPR/Cas9-based screens further prompted us to perform genome-wide profiling of vulnerabilities to ATR inhibition (Wang et al., 2019). These unbiased profiling of genes when lost may affect sensitivity to ATR inhibition is useful for at least two purposes. First, these studies allow the identification of new genetic alternations and biomarkers that endow hypersensitivity of tumors to ATR inhibition, which can be exploited in clinic. Second, these studies also reveal novel genes that have functional and/or genetic interaction with ATR pathway.

In this study, we will focus on a previously uncharacterized gene C17orf53, whose loss led to increased sensitivity to ATR inhibition in our previous screen (Wang et al., 2019). To date, human C17orf53 was only reported in few studies and the functions of C17orf53 remain largely unknown. A previous study indicated a potential relationship between C17orf53 SNPs and bone density (Styrkarsdottir et al., 2009), while another recent study suggested that C17orf53 is a G2/M specific gene (Giotti et al., 2018).

Here, we showed that C17orf53 is conserved in vertebrate and its expression associates with cell proliferation. Loss of C17orf53 induced aberrant DNA replication and pronounced ICL repair defect. Moreover, C17orf53 functions independent of FA pathway in ICL repair. Its depletion impaired the loading of late HR factors. Furthermore, we showed that C17orf53 is ssDNA- and RPA-binding protein. Collectively our data suggest that C17orf53 may act in a new pathway that promote ICL repair independent of FA pathway.

## Results

### C17orf53 loss sensitizes cells to ATR inhibition

We recently reported genome-wide CRISPR screens for profiling genes that affect cellular response to ATR inhibition (Wang et al., 2019) (**Figure 1A**). In this study, many genes involved in DNA replication or replication-related DNA repair pathways were enriched, which include FA, HR and NER pathways (**Figure 1B**). We found 24 candidate genes correlated robustly with response to ATR inhibition, with 18 genes whose loss induced hypersensitivity to ATRi and 7 genes whose loss induced resistance to ATRi (**Figure 1C**). Some of these candidate genes have been already reported by us and others (Cotta-Ramusino et al., 2011; Lee et al., 2007; Ruiz et al., 2016; Wang et al., 2019).

**Figure 1.**
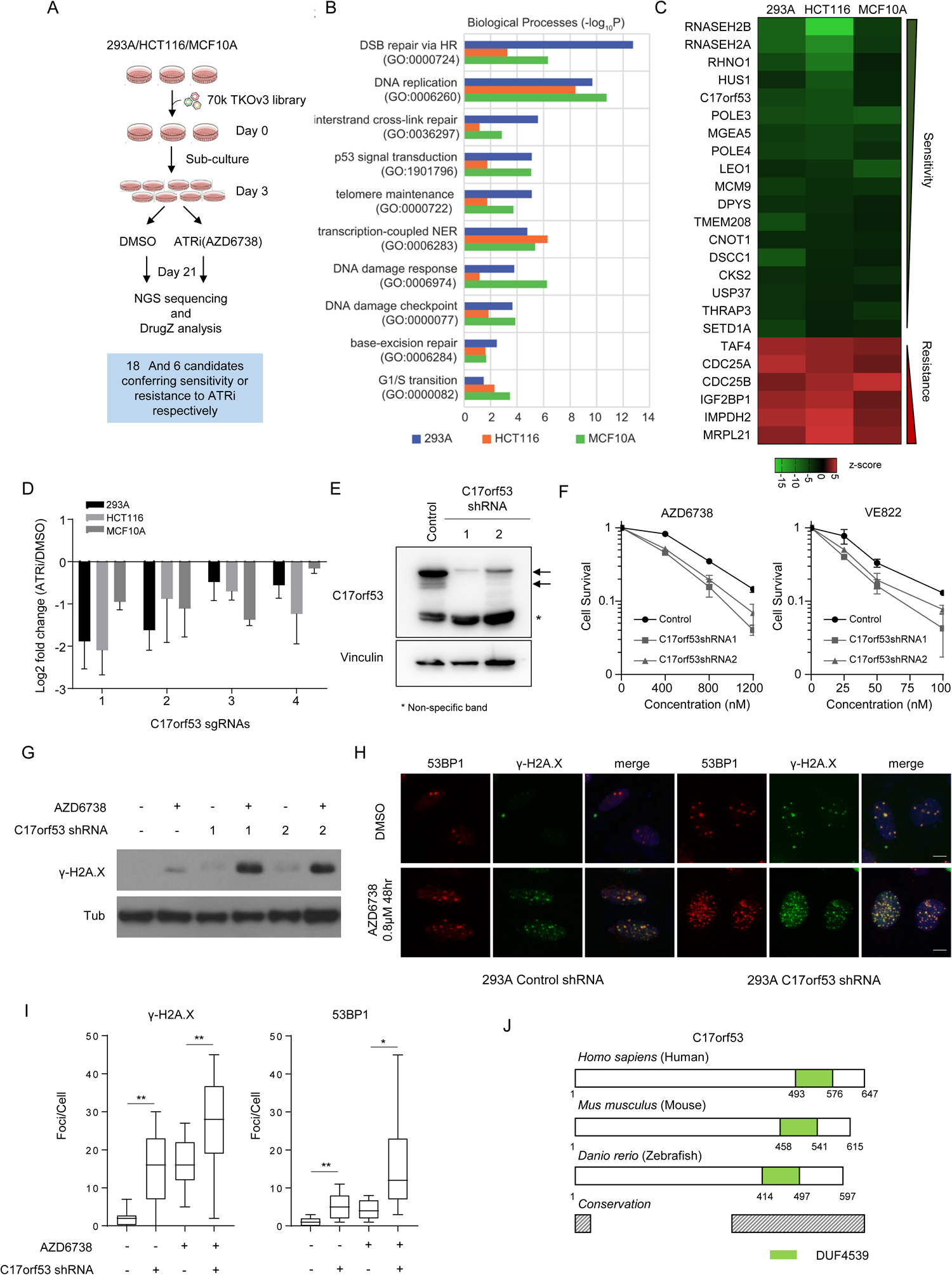
CRISPR screens identified C17orf53 as a determinant of ATRi sensitivity. **(A)** Schematic representation of the workflow for CRISPR screens performed in 293A, HCT116, and MCF10A cells. **(B)** Gene ontology (GO) analysis of the synthetic lethality genes identified in the CRISPR screens. The top10 GO terms significantly enriched were listed here. Different color represents different cell line. Blue: 293A, Orange: HCT116, Green: MCF10A**. (C)** Visualization of the high-confidence genes in these screens. The drugZ scores are displayed as a heatmap summarizing the common genes uncovered in all three screens (FDR<0.01) which show synthetic lethality (score<0) or synthetic survival (score>0) with ATR inhibition. **(D)** Fold change ratios of sgRNAs targeting C17orf53 in 3 cell lines comparing ATRi treated group with DMSO treated group. Mean±SD, n=2. (**E)** Immunoblot of wild-type and C17orf53 knock-down 293A cells. **(F)** Clonogeneic survival assay of cells treated with ATR inhibitors AZD6738 or VE-822. Mean±SD, n=3. **(G)** Wild type and C17orf53 knock-down 293A cells were treated with AZD6738 (0.8µM, 48hr). Cells were harvested and lysates were analyzed by immunoblotting with γ-H2A.X and tubulin antibodies. **(H**) Wild type and C17orf53 knock-down 293A cells were treated with DMSO or AZD6738 (0.8µM, 48hr). Cells were fixed and processed for γ-H2A.X and 53BP1 immunofluorescence staining. The scale bar represents 1 µm. **(I)** Quantification of γ-H2A.X and 53BP1 foci in different groups in H). n=100 cells, *P<0.05, **P<0.01, student t-test. **(J)** Schematic diagram of C17orf53 proteins from human to zebrafish.

We focused on an uncharacterized gene *C17orf53*, loss of which induced hypersensitivity to ATR inhibition, since significant dropout of sgRNAs targeting this gene was noted in ATR inhibitor treated group (**Figure 1D**). We first confirmed these screening results with a different gene depletion method. Using shRNAs, we efficiently knocked down *C17orf53* in 293A cells, which were validated in both mRNA and protein levels (**Figure 1E and Supplementary Figure 1A**). Cell survival assays showed that the *C17orf53* knock-down cells were sensitive to treatment with two different ATR inhibitors (**Figure 1F and Supplementary Figure 1B**), further confirming that *C17orf53* is a determinant of ATRi sensitivity.

On molecular levels, ATR inhibition induced dramatic increase of γ-H2A.X in *C17orf53* knock-down cells (**Figure 1G**). Moreover, γ-H2A.X and 53BP1 foci were also significantly increased in *C17orf53* knock-down cells upon ATR inhibitor treatment (**Figure 1H, I**), indicating more DSBs were generated due to depletion of *C17orf53*. All these results suggest a possible function of C17orf53 in genome maintenance. Furthermore, as genes co-essential with ATR inhibition usually participate in DNA replication or repair (Rundle et al., 2017; Saldivar et al., 2017; Wang et al., 2019), our data indicate that C17orf53 may function in these processes.

### C17orf53 is a conserved gene involved in cell proliferation

C17orf53 is highly conserved in vertebrates, which is schematically depicted in Figure 1J (please also see **Supplementary Figure 1C** for sequence alignment). While the function of C17orf53 remains unknown, a previous study indicated *C17orf53* SNP correlates with BMD disease (Styrkarsdottir et al., 2009). Tissue expression profiling showed *C17orf53* is highly expressed in bone marrow and testis (**Supplementary Figure 2A**), both of which are highly proliferative adult tissues, indicating a potential correlation between C17orf53 expression and cell proliferation.

To examine *C17orf53* expression in tumors (hyper-proliferating cells) and normal tissues (with limited proliferating cells), we first screened *C17orf53* expression profiles in a broad number of cancers (**Figure 2A**). We found *C17orf53* was significantly up-regulated in most cancer types comparing with the relevant normal tissue (**Supplementary Figure 2B**). Moreover, we found the expression of *C17orf53* correlated robustly with numerous genes involved in DNA replication and replication-associated DNA repair processes such as HR and FA pathways in multiple datasets (**Figure 2B, C, D and Supplementary Figure 2C**). These data further implicated the relationship between *C17orf53* expression and cell proliferation

**Figure 2.**
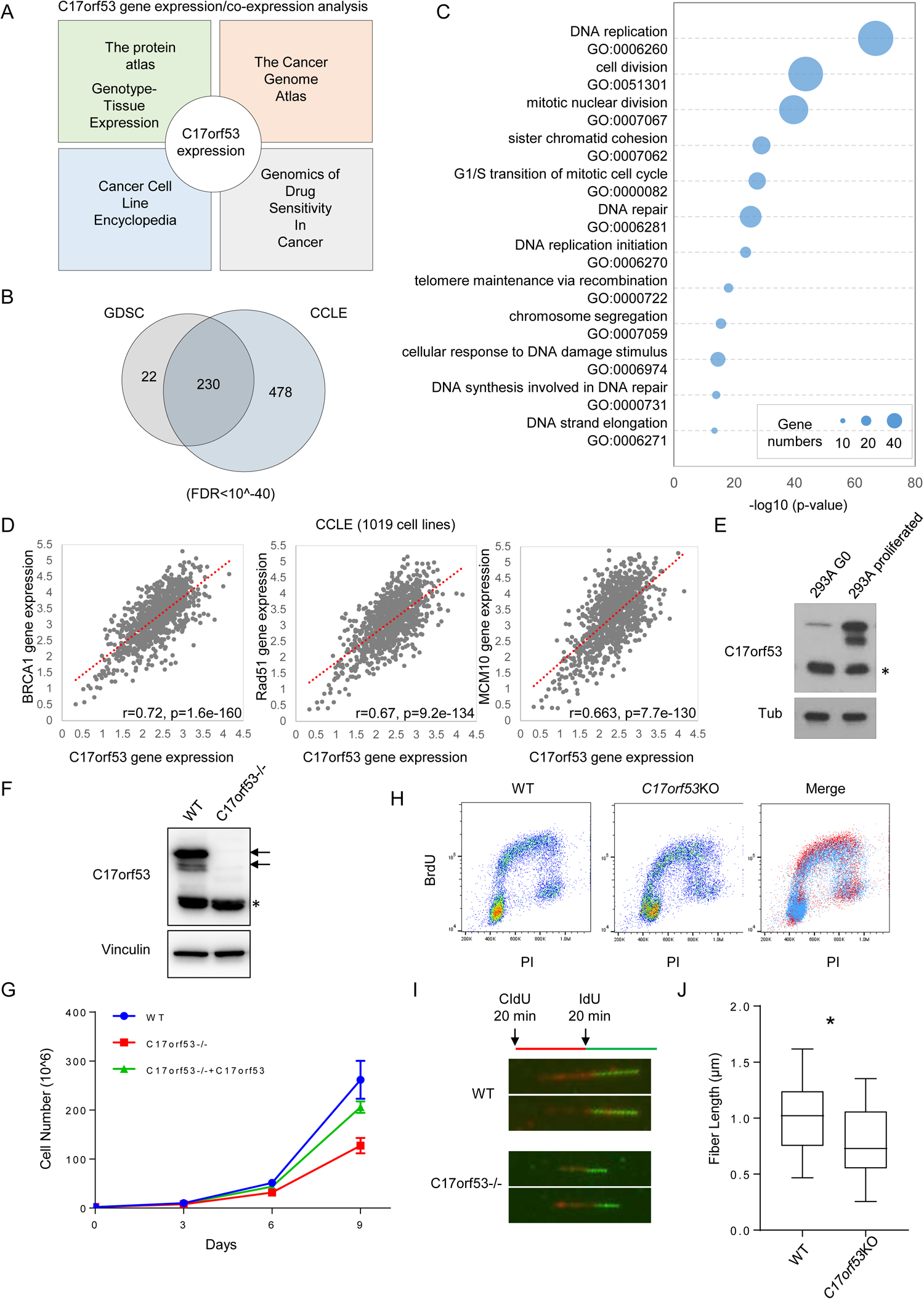
C17orf53 expression correlates with expression of DNA replication and HR genes. **(A)** Overview of *C17orf53* gene co-expression analysis. **(B)** Venn diagram of high-confidence co-expressed genes with *C17orf53* derived from two datasets. (p<10e-40, CCLE, GDSC). **(C)** GO terms significantly (p<10e-10) enriched among genes common to the two datasets in B). **(D)** *C17orf53* expression correlates with BRCA1, RAD51 and MCM10 in 1019 cell lines collected in CCLE. **(E)** Comparison of 293A cells that were contact inhibited (G0) and 293A cells that were 12hrs after released from contact inhibition (proliferating). Total cell lysates were immunoblotted with the indicated antibodies. * marks the non-specific band. **(F)** Immunoblot of wild-type and C17orf53KO cells. **(G)** Growth curves of indicated cells. Different cells were plated in 6-well plates, passaged and counted every 3 days. (**H)** BrdU incorporation assay. Growing wild-type and C17orf53KO 293A cells were incubated in BrdU for 30mins, fixed and stained with BrdU antibody and PI. (**I)** DNA fiber lengths in wild-type and C17orf53KO 293A cells. **(J)** Quantification of DNA fiber lengths in I). The box plots show twenty-fifth to seventy-fifth percentiles with lines indicating the median.

In 293A cells, *C17orf53* is highly expressed in proliferating cells compared with that in quiescent cells (**Figure 2E).** A recent study suggested that *C17orf53* is a highly expressed gene in S/G2 cell cycle phase (Giotti et al., 2018). However, in our experiment, the C17orf53 protein level did not change significantly during cell cycle (**Supplementary Figure 2E**). These findings indicate that *C17orf53* is a cell proliferation related gene but its protein level is not significantly regulated during cell cycle.

It is also of note that the *C17orf53* knock-down cells showed retarded cell growth and clonogenic ability, suggesting that *C17orf53* is necessary for optimal cell proliferation. To confirm this result, we generated *C17orf53* knock-out cells using CRISPR/Cas9 gene editing technology, which was validated by immunoblotting and genome sequencing (**Figure 2F and Supplementary Figure 2F**). Consistent with those observed in *C17orf53* knock-down cells, the *C17orf53*-KO cells also showed slow cell growth and decreased clonogenic ability which could be rescued by reconstruction with ectopically expressed C17orf53 (**Figure 2G**).

The nice correlation of *C17orf53* with numerous genes required for faithful DNA replication (**Figure 2B, C, D and Supplementary Figure 2C**) implied a similar role for C17orf53. Thus, we checked DNA replication in *C17orf53* KO cells. As shown in **Figure 2H**, although *C17orf53* KO cells showed similar cell cycle distribution as that of WT cells, BrdU incorporation was reduced in *C17orf53* KO cells. DNA fiber assay further confirmed slowed replication fork progression in *C17orf53* KO cells (**Figure 2I, J**). These findings suggest loss of *C17orf53* may affect DNA replication, which may be one of the reasons that these cells are hypersensitive to ATR inhibition.

### C17orf53 depleted cells show profound ICL repair defect

To gain more insight into C17orf53 functions, we tested the sensitivity of *C17orf53* KO cells to treatment with a variety of DNA-damaging agents. We further confirmed that loss of *C17orf53* sensitized cells to ATR inhibition and Chk1 inhibition (**Figure 3A and Supplementary Figure 3A**). Besides, *C17orf53* KO cells showed hypersensitive to MMC treatment, but not to other agents such as IR, UV, HU and CPT (**Figure 3A and Supplementary Figure 3A**). Compared to WT cells, *C17orf53* KO cells showed significant G2/M arrest upon MMC treatment (**Figure 3B**). Moreover, MMC treatment induced exacerbated DSB checkpoint activation as indicated by the dramatic increase of pRPA2 in *C17orf53* KO cells, while neither HU nor CPT treatment could induce such dramatic change in *C17orf53* KO cells (**Figure 3C**). This result was further confirmed with treatment of different doses of MMC, which showed pPRA2 and γH2A.X levels were significantly increased in *C17orf53* KO cells upon MMC treatment (**Figure 3D**). We also generated *C17orf53*KO U2OS cells. Similar to *C17orf53*KO 293A cells, MMC treatment could induce dramatic G2/M arrest and significant pRPA2 increase in *C17orf53* depleted U2OS cells (**Supplementary Figure 3B, C**). Moreover, the luciferase based ICL repair reporter assay showed that, comparing to WT cells, *C17orf53* KO cells showed decreased ability to repair the crosslink in the reporter plasmid while ectopic expression of C17orf53 could partially rescue it (**Supplementary 3D, E**). These results above showed that C17orf53 depleted cells harbored profound defects in response to ICL-inducing agent, suggesting a function of C17orf53 in ICL repair.

**Figure 3.**
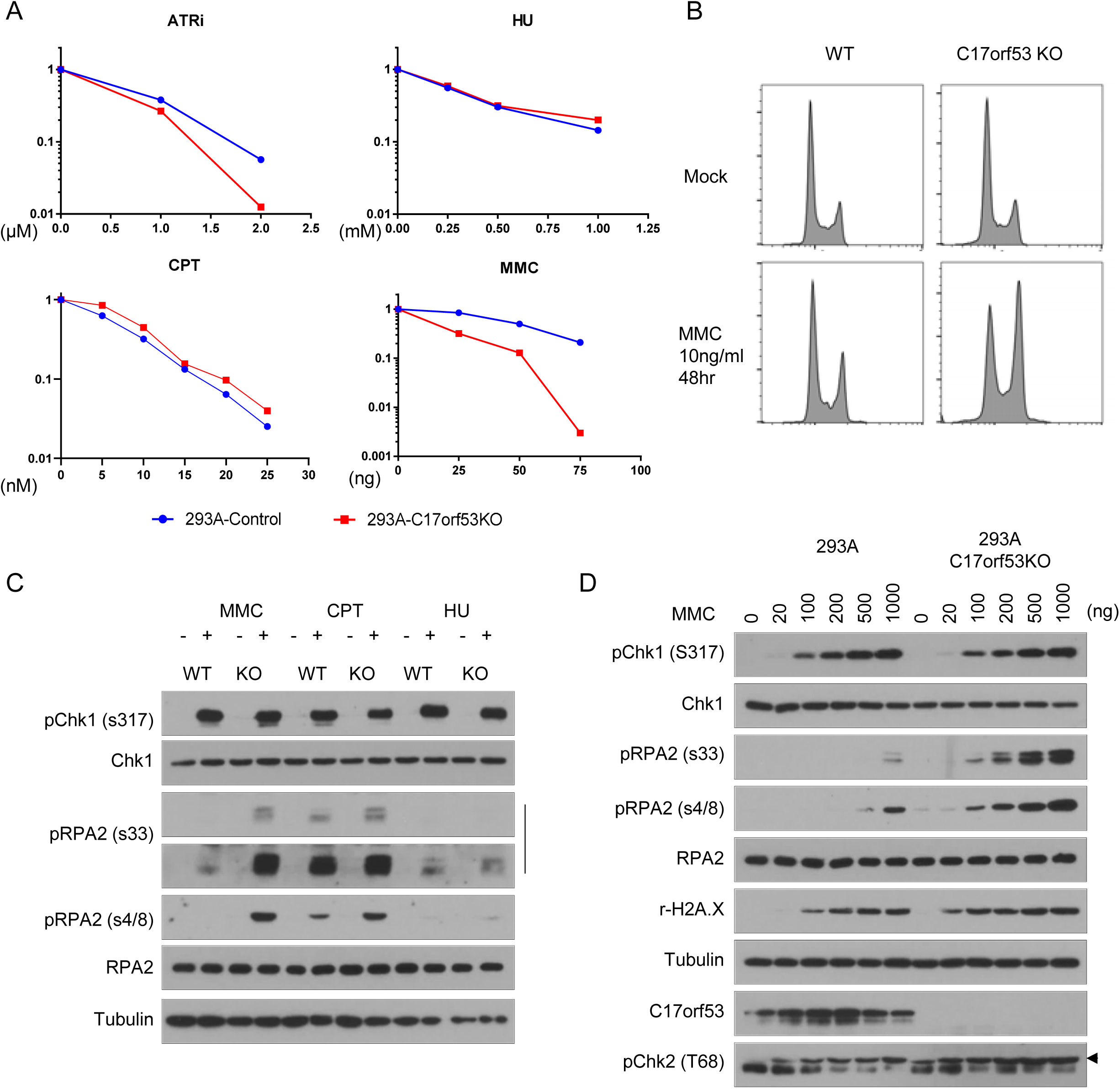
C17orf53 deficient cells show defects in ICL repair. **(A)** Clonogenic survival analysis of wild-type and *C17orf53*KO 293A cells exposed to ATRi (AZD6738), HU, CPT and MMC. Mean and SD of two independent experiments. **(B)** Cells were treated with DMSO or MMC (10 ng/ml) for 48 hrs before ethanol fixation, propidium iodide staining, and fluorescence-activated cell sorting (FACS) analysis**. (C)** Wild-type or *C17orf53*KO 293A cells were treated with MMC (500 ng/ml), HU (2 mM) or CPT (200 nM) for 8 hrs and then lysed. The total cell lysates were immunoblotted with the indicated antibodies. **(D)** Wild-type or *C17orf53*KO 293A cells were treated with MMC of different doses for 8 hrs and then lysed. The total cell lysates were immunoblotted with the indicated antibodies.

### C17orf53 functions in the late stage of ICL repair that is downstream of FANCD2/RPA/RAD51

ICL repair is a complex process that requires the action of multiple DNA repair pathways [25-30]. To address the function of C17orf53 in this process, we treated WT or *C17orf53* KO cells with MMC for 1 hour, released them from MMC, and then monitored DNA repair and DNA damage markers in these cells over time. We found that pRPA2 level was significantly increased in *C17orf53* KO cells at any time point (**Figure 4A**). We also noticed that, in *C17orf53* KO cells, γH2A.X level kept increasing (**Figure 4A**), indicating an accumulation of DNA damage, which agrees with ICL repair defect caused by *C17orf53* depletion. Consistent with the observations above, *C17orf53* KO cells showed significant G2/M arrest even 48 hrs after MMC treatment (**Supplementary 4A**).

**Figure 4.**
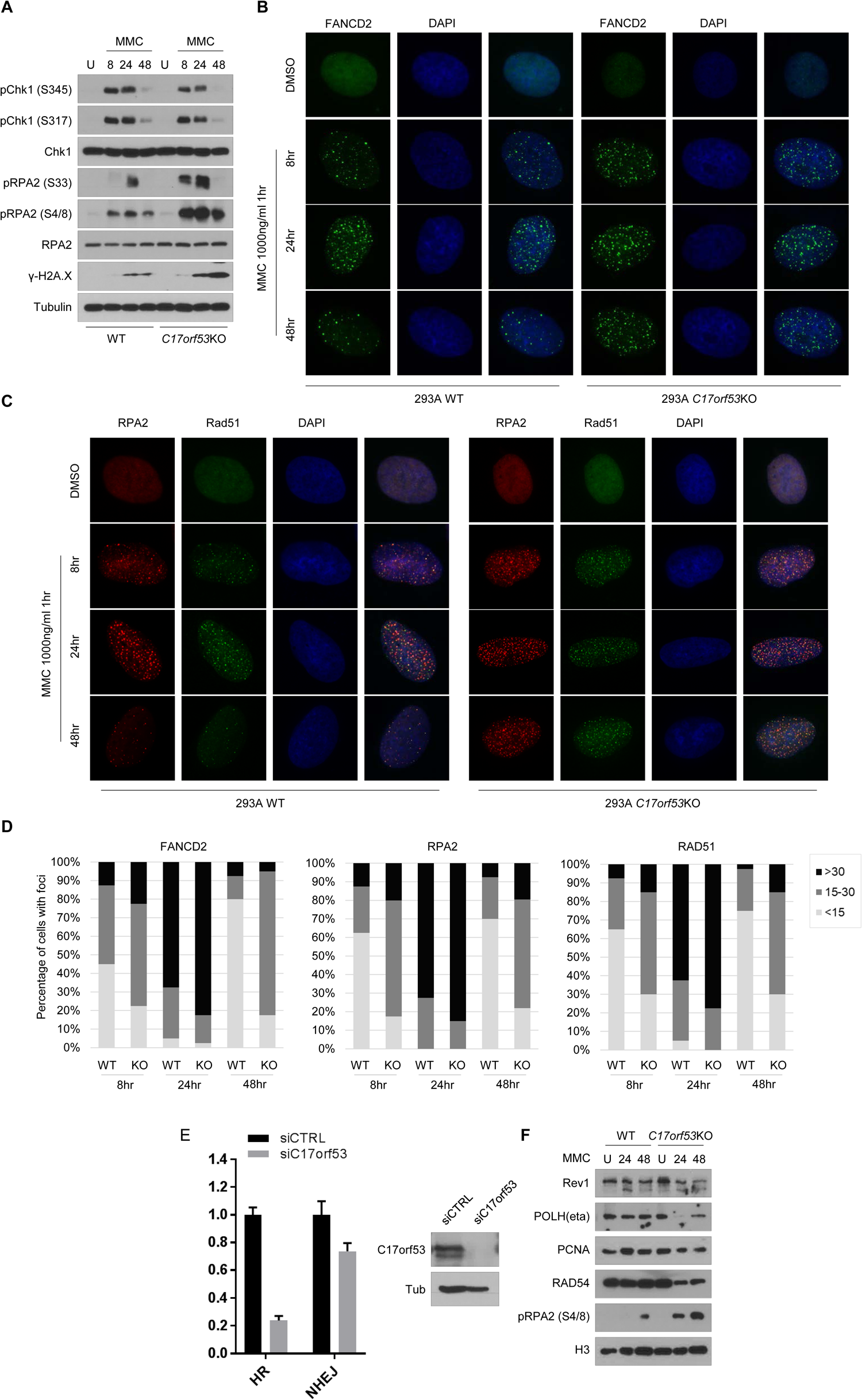
C17orf53 participates in ICL repair through affecting HR repair. **(A)** Wild-type cells and *C17orf53* KO cells were treated with DMSO or MMC (1000 ng/ml, 1 hr). The released cells at different time point were harvested and immunoblotted with the indicated antibodies. **(B)** MMC-induced FANCD2 foci in wild-type and C17orf53 KO 293A cells. The indicated cells were left untreated or treated with MMC (1000 ng/ml, 1hrs). The released cells at different time point were fixed and immunostained with FANCD2 antibody. **(C)** MMC-induced RPA2 and RAD51 foci in wild-type and C17orf53KO 293A cells. The indicated cells were left untreated or treated with MMC (1000 ng/ml, 1hr). The released cells at different time point were fixed and immunostained with RPA2 and RAD51 antibodies. **(D)** Quantification of RPA2, RAD51 and FANCD2 foci in B) and C). n=40 cells. **(E)** Loss of C17orf53 results in decreased HR-directed DNA repair. U2OS-DR-GFP or U2OS-NHEJ-GFP cells were transfected with control siRNA or C17orf53 siRNA, followed by transfection of I-SceI. 48hr after transfection, cells were harvested and assayed for GFP expression by FACS. Mean±SD, n=2. The knockdown efficiency were validated by immunoblotting with C17orf53 antibody. **(F)** Cells in A) were harvested, fractionated and analyzed by immunoblotting with the indicated antibodies.

We also monitored repair kinetics through enumeration of FANCD2, RPA, RAD51 foci formation. Immunostaining results showed, in *C17orf53* KO cells, MMC treatment induced more FANCD2 foci at the early time point (8hrs), which also persisted much longer (48hrs) than those in WT cells (**Figure 4B**), suggesting that while the recruitment of FANCD2 was not affected by *C17orf53* depletion, the ICL repair was impaired. Consistently, the monoubiquitination of FANCD2 and FANCI were not changed in *C17orf53* KO cells comparing to WT cells. Similar to FANCD2 foci, RPA2 and RAD51 foci increased significantly at early time point (8hrs) and persisted much longer (48hrs) (**Figure 4C**), which again suggested that DNA resection indicated by RPA foci and the recruitment of RAD51 were not affected in *C17orf53*KO cells.

Together, these data suggest that C17orf53 functions in the late stage of ICL repair. We then used well-established reporter assays to monitor HR- or NHEJ-mediated capacity. We knocked down *C17orf53* using siRNA in U2OS-DR-GFP cells and found that C17orf53 depletion led to a pronounced reduction in HR repair efficiency (**Figure 4E**). Interestingly, overall NHEJ repair efficiency was also impaired, albeit not as severe as HR repair, in these cells. These results suggest that defective ICL repair observed in *C17orf53* depleted cells may be at least in part due to defective HR repair measured by these reporter assays. However, the underlying mechanisms may be distinctively different, since *C17orf53* KO cells did not show significant sensitive to IR or CPT treatment.

To gain further insight into the mechanism underlying ICL repair defect observed in C17orf53 depleted cells, we examined the chromatin loading of several repair factors and showed that the chromatin loading of Rev1, PolH, PCNA and RAD54 were compromised in *C17orf53* KO cells (**Figure 4F**). Precisely how C17orf53 influences the loading of these repair factors remains to be elucidated.

### CRISPR screens with DDR library in WT and C17orf53KO cells reveal genetic interaction between C17orf53 and FA/HR pathway

We decided to take advantage of CRISPR screens to elucidate functional and/or genetic interaction between *C17orf53* and other DDR genes. We created a targeted DDR library including 358 genes with known or suspected functions in DDR. As described in **Figure 5A**, WT cells or *C17orf53* KO cells were infected with lentivirus encoding gRNAs targeting these DDR genes or lentivirus for the whole genome TKOv3 library and selected by puromycin. After 20 population doubling, cell pellets in each group were collected and the relevant changes of sgRNAs were determined by next generation sequencing of PCR-amplified samples. The sequencing results were analyzed with MAGeCK.

**Figure 5.**
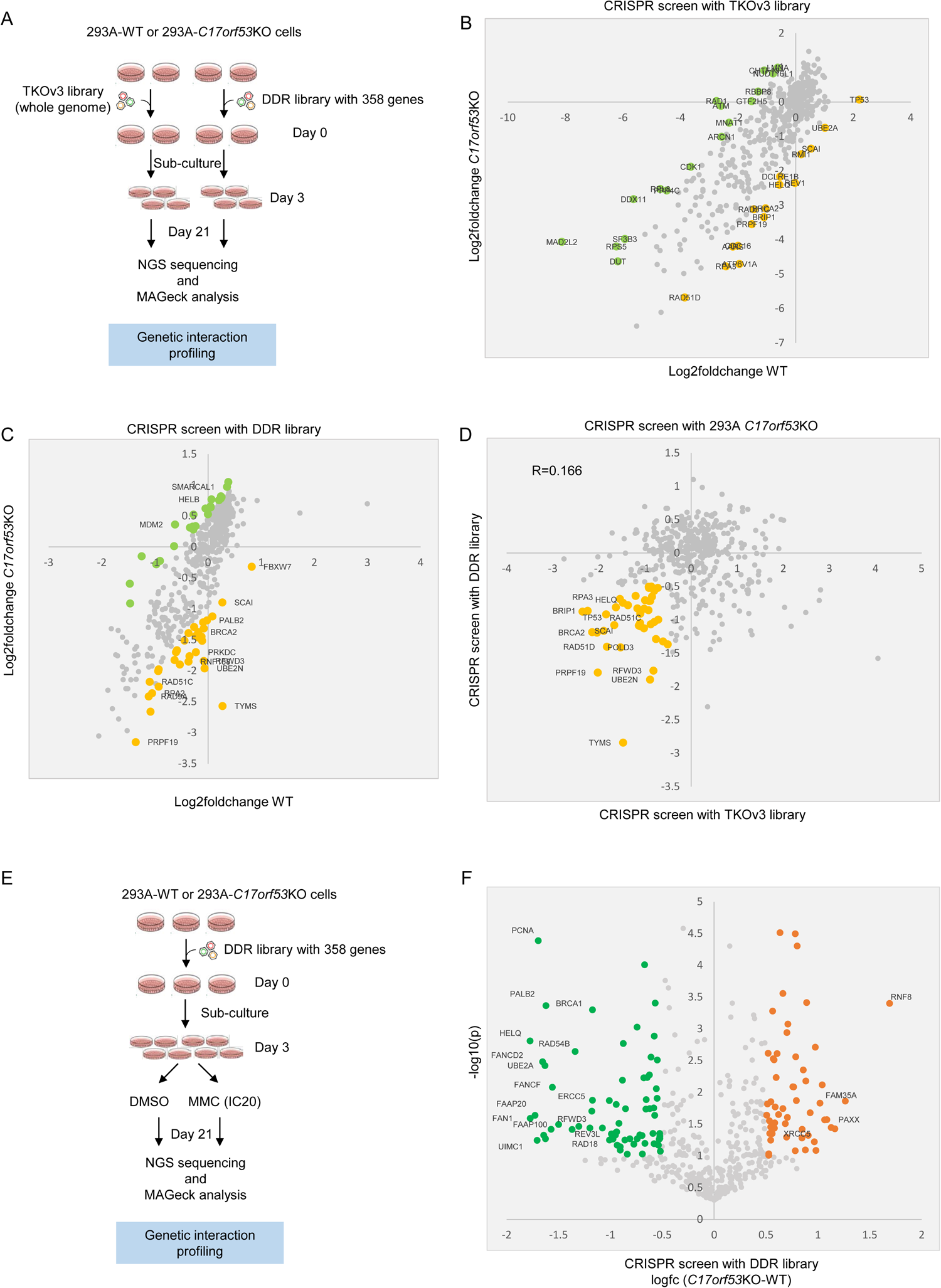
Genetic interaction profiling by CRISPR screens. **(A)** Schematic representation of the workflow for CRISPR synthetic lethality screens performed in wild-type and *C17orf53* KO 293A cells with TKOv3 whole-genome library and DNA damage response (DDR) library. **(B-C)** Results from the CRISPR TKOv3 whole genome (B) and CRISPR DDR (C) screens, plotted as the log2 fold-change in *C17orf53*KO cells against the log2 fold-change in wild-type cells**. (D)** Comparison of genetic interaction (GI) score (log2 fold-change for each gene in *C17orf53*KO versus wild-type cells) in TKOv3 screen and DDR screen. Genes that have GI>1 are color-coded as orange for relative drop-out or green for relative enrichment. **(E)** Schematic representation of the workflow for CRISPR screens performed in wild-type and *C17orf53* KO 293A cells with DDR library upon MMC treatment. **(F)** Volcano plots for CRISPR screens exposed to MMC treatment in wild-type and *C17orf53*KO cells. Significant genes are color-coded as orange for relative drop-out or green for relative enrichment.

From the analysis of our screens, several themes emerged. We first noticed that depletion of TYMS, RFWD3, SCAI, HELQ, UBE2N genes inhibited growth in *C17orf53* KO cells more than they did in WT cells, yielding a KO/WT ratio that is negative on a log2 scale; this ratio can also be negative for genes that promote growth of WT cell more than *C17orf53*KO cells (**Figure 5 B, C, D**). Several of these genes participate in DNA replication and RPA modification, suggesting the physiological function of C17orf53 may be involved in these processes, which is also consistent with our pervious observations.

We also performed experiments using DDR library in WT cells or *C17orf53* KO cells treated with DMSO or MMC (**Figure 5E**). When under MMC treatment, WT cells appeared to be more reliant on FA/HR/TLS pathways though loss of FA/HR/TLS pathways were also toxic to both WT and *C17orf53* KO cells, while NHEJ pathway components were more essential for the survival of *C17orf53* KO cells (**Figure 5F**). In the other words, these epistasis genetic screens under MMC treatment showed that loss of *C17orf53* did not exacerbate ICL repair defects when FA/HR/TLS pathways were impaired, suggesting C17orf53 and FA/HR/TLS may operate together to promote ICL repair. In the absence of C17orf53, cells with compromised ICL repair appears to accumulate more DSBs as we demonstrated above and therefore these cells have an increased need for DSB/NHEJ repair for their survival.

### C17orf53 contains a ssDNA-binding domain-DUF4539, which is essential for its function

To clarify the molecular basis of C17orf53 function, we decided to further characterize C17orf53. The highly conserved C-terminus of C17orf53 contains an unknown domain, DUF4539 (**Figure 6A).** Bioinformatics analysis revealed a putative OB fold within DUF4539, suggesting that this domain may be able to bind to nucleotides (**Supplementary Figure 5A, B**). EMSA assays using purified proteins showed that the C-terminus of C17orf53 containing the DUF4539 domain could preferentially bind to ssDNA, but not dsDNA (**Figure 6B**). Deletion of DUF4539 abolished the ssDNA-binding ability of the C-terminus of DUF4539 (**Figure 6B, C**). Reconstruction experiments in *C17orf53* KO cells showed that ectopic expression of C17orf53-delDUF4539 induced more severe G2/M arrest upon MMC treatment **(Figure 6D)**, suggesting ssDNA binding is essential for the function of C17orf53.

**Figure 6.**
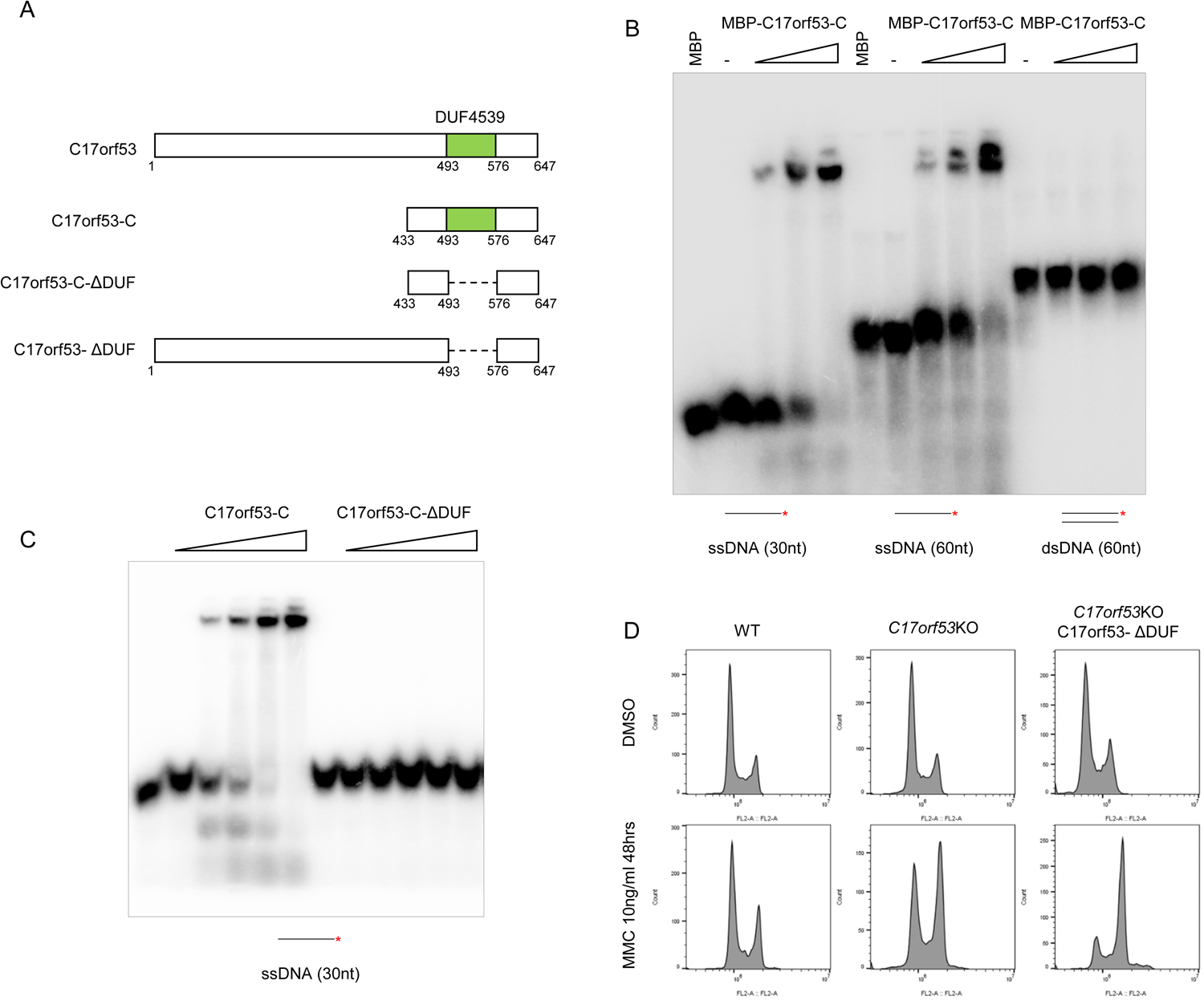
C17orf53 prefers to bind to ssDNA through DUF4539 domain. **(A)** Schematic diagram of C17orf53 indicating the location of DUF4539 domain and the truncation fragments tested in B) and C). **(B)** Electrophoretic mobility shift analysis of the indicated DNA ligands incubated with MBP-C17orf53-C**. (C)** Electrophoretic mobility shift analysis of the indicated DNA ligands incubated with MBP-C17orf53-C and MBP-C17orf53-CΔDUF4539 (i.e. C17orf53-CΔDUF). **(D)** Indicated cells were treated with DMSO or MMC (10 ng/ml) for 48 hrs before ethanol fixation, propidium iodide staining, and fluorescence-activated cell sorting (FACS) analysis.

### C17orf53 is a RPA binding protein

We also performed TAP/MS to search for C17orf53-binding partners. As shown in **Figure 7A**, the C17orf53 interactome revealed enrichment of RPA complex. We conducted CoIP experiments and pull-down assays to validate the interaction between RPA complex and C17orf53. The CoIP experiments showed that C17orf53 could bind strongly to RPA1 and RPA2 (**Figure 7B**). Further *in vitro* pull-down assays demonstrated that RPA1 and RPA2 could bind directly to C17orf53 (**Figure 7C**). Further mapping results showed that C17orf53 binds to RPA1 and RPA2 through two separate RBMs at respectively N- and C-terminus of C17orf53 (**Figure 7 D, E, F, G**).

**Figure 7.**
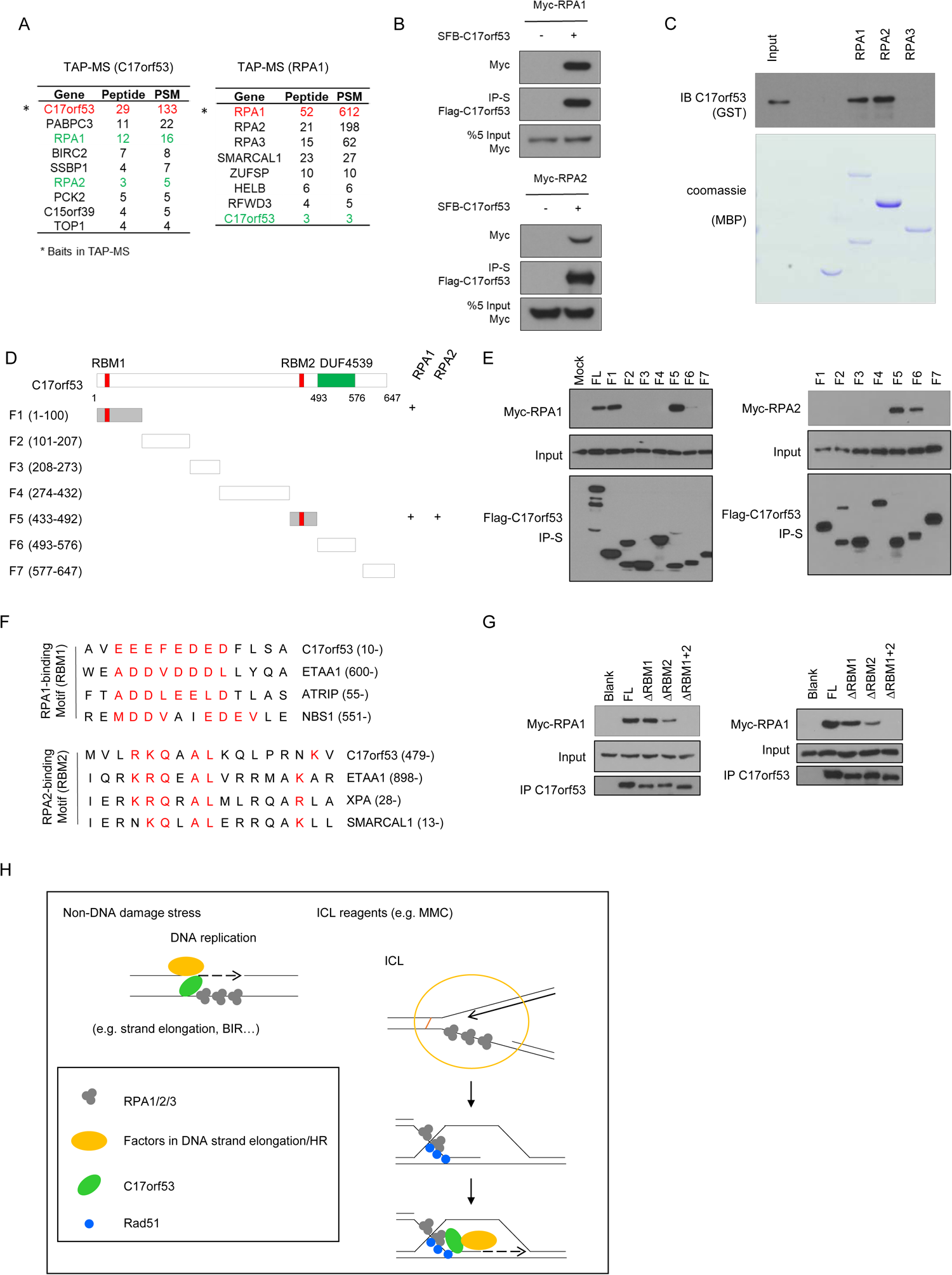
C17orf53 is a RPA binding protein. **(A)** 293T cells stably expressing SFB-C17orf53 or SFB-RPA1 were used for TAP of the protein complexes. Tables are summaries of proteins identified by mass spectrometry analysis. Letters in Red indicate the bait proteins. **(B)** C17orf53 interacts with RPA1 and RPA2. 293T cells were transiently transfected with indicated plasmids. Cell lysates were immunoprecipitated with S beads, and immunoblot was performed with the indicated antibodies. **(C)** Direct binding between recombinant GST-C17orf53 and MBP-tagged RPA1 and RPA2. Upper panel: C17orf53 was detected by immunoblotting. Lower panel: purified proteins were visualized by Commassie staining**. (D)** Schematic diagram of C17orf53 indicating the truncation fragments tested in E) and G). **(E)** 293T cells were transiently transfected with indicated plasmids. Cell lysates were immunoprecipitated with S beads, and immunoblot was performed with the indicated antibodies. **(F)** Alignments of RPA1- and RPA2-binding motifs (RBM1 and RBM2) across multiple proteins. **(G)** 293T cells were transiently transfected with indicated plasmids. Cell lysates were immunoprecipitated with S beads, and immunoblot was performed with the indicated antibodies. **(H)** The proposed function model of C17or53. Without DNA damage, C17orf53 works for DNA replication. When facing with MMC-induced ICL, C17orf53 functions at the late stage to sufficiently recruit late HR factors.

## Discussion

In this study, we focused on a previously uncharacterized gene, *C17orf53*, which we identified as a gene that promotes viability especially in cells treated ATR inhibitor. We validated that loss of C17orf53 sensitized cells to ATR inhibition. We showed that *C17orf53* expression correlates with cell proliferation. C17orf53 may be important for cell proliferation due to its possible function in DNA replication. Interestingly, *C17orf53* depleted cells showed profound ICL repair defect. Further experiments revealed that C17orf53 functions at the late stage of ICL repair. Moreover, loss of *C17orf53* led to aberrant HR repair, probably through impairing loading of several repair factors. In parallel, we performed functional CRISPR screens and revealed that C17orf53 may participate in ICL repair together with FA/HR proteins. On molecular levels, C17orf53 is a ssDNA-binding and RPA-binding protein, both of which may be essential for its function in ICL repair. Taken together, our study revealed a new protein that functions in ICL repair. Further investigation of this process may reveal mechanistically how C17orf53 participates in ICL repair independent of FA and HR, the other two repair pathways known to be involved in ICL repair.

The high *C17orf53* expression across many cancer types and the correlation of C17orf53 expression with poor diagnosis suggest a possibility that C17orf53 could be a reliable marker to discriminate pre-cancerous cells from normal cells. It is also of note that C17orf53 is a conserved protein, but seems to appear exclusively in vertebrate. Considering reduced DNA replication observed in cells with loss of *C17orf53*, it seems that the increased genome complexity may require more sophisticated regulation and thus the functions of C17orf53 during replication and repair. Nevertheless, the detailed mechanisms underlying the functions of C17orf53 in DNA replication and ICL repair remain to be further elucidated.

## AUTHOR CONTRIBUTIONS

C.W. and J.C. conceived the project. C.W., Z.C., S.D., M.T., L.N., H.Z. and X.F. performed the experiments. M.M. provided technical support for the screen work. R.W., X.S. and L.L. provided instruction about the study of ICL repair. C.W. and T.H. analyzed the deep-sequencing results. C.W. and J.C. wrote the manuscript with input from all authors.

## ACKNOWLEDGEMENT

We thank the members of Dr. Junjie Chen’s lab for their kind help. U2OS DR-GFP and NHEJ-GFP cells were kindly provided by Dr. Albert C. Koong at the University of Texas MD Anderson Cancer Center. This work was supported in part by CPRIT (RP160667) and NIH grants (CA157448, CA193124, CA210929, CA216911, and CA216437) to J.C. and MD Anderson’s NIH Cancer Center Support Grant (CA016672).

## Methods and Materials

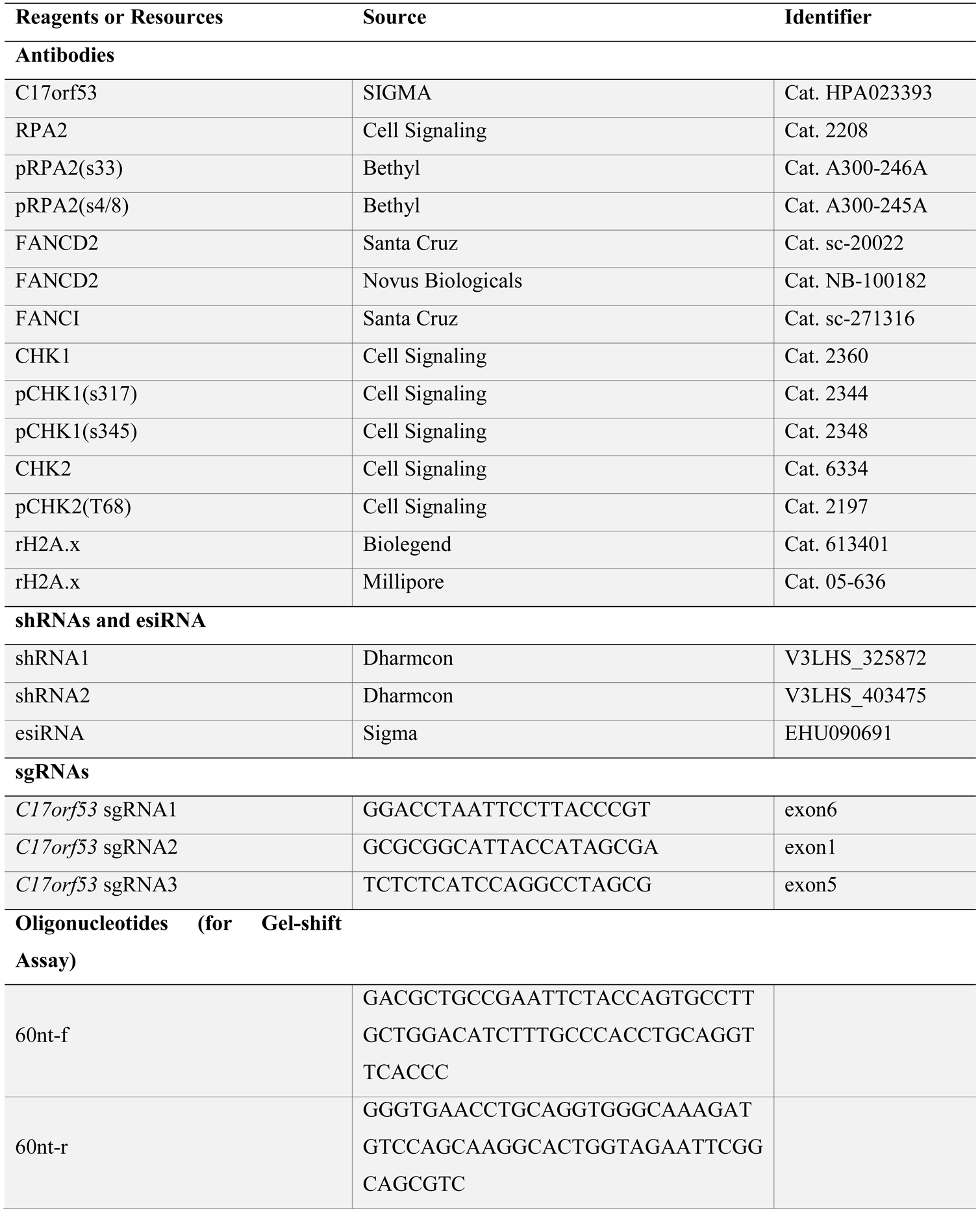
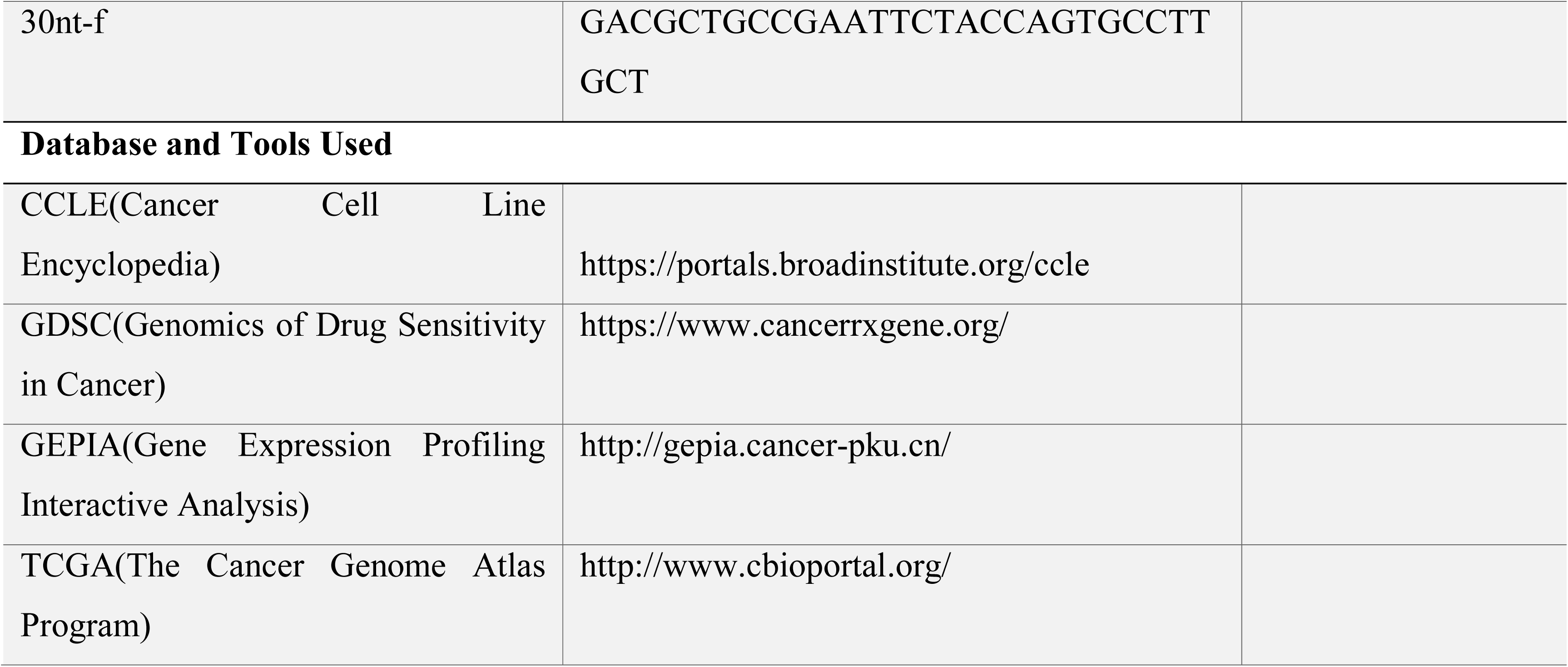

### Cell Culture

293A, HCT116, MCF10A, and U2OS cells were obtained from ATCC. Dulbecco’s modified Eagle medium (DMEM) with 10% fetal calf serum was used to culture 293A, HCT116, and U2OS cells. MCF10A cells were cultured in DMEM/F12 Ham’s mixture supplemented with 5% equine serum (Gemini Bio), 20 ng/mL epidermal growth factor (Sigma), 10 μg/mL insulin (Sigma), 0.5 mg/mL hydrocortisone (Sigma), 100 ng/mL cholera toxin (Sigma), 100 units/mL penicillin, and 100 μg/mL streptomycin.

### Viability Assay

For cell growth assays, cells were seeded in 6-well plates (10^4^ per well) and passaged every 3 days until cell numbers were determined. For clonogenic assays, cells were seeded in 6-well plates (400 cells per well) and continuously exposed to chemicals treatment for 14 days, beginning 24 h after seeding. Cells were fixed with 10% trichloroacetic acid and stained with sulphorhodamine B (Sigma-Aldrich). Colonies were counted manually. All cell survival assays were performed at least in triplicate.

### Cell Cycle Analysis by Flow Cytometry

Cells were incubated with 10µM 5-bromo-2-deoxyuridin (BrdU) for 30min before fixation with 70% ethanol. BrdU was detected with BrdU antibody. Propidium iodide (PI) was used to measure DNA content. Data were collected with BD C6 (Becton Dickinson) and analyzed with Flowjo.

### Immunofluorescence staining Analysis

Cells were grown on coverslips for 24 h before treatment. After the indicated treatment, cells were fixed in 4% paraformaldehyde and permeablized with phosphate-buffered saline (PBS) with 0.5% Triton X-100. Then, cells were incubated with primary antibodies diluted in PBS with 0.05% Triton X-100 and 1% BSA (PBST-BSA) for 1h at room temperature. After 3 washes with PBS, fluorescently labeled secondary antibodies in PBST-BSA were added for 1 h. Cells were then washed in PBS with Hoechst stain (1:10,000). Slides were imaged at 40× on a Leica microscope.

### Generation of knock-out cells

pLentiCRISPRv2 was used to generate KO cells. Cells were transiently transfected with the indicated plasmids and selected using puromycin (2 mg/mL). Single cells were then plated into 96-well plates. After 10 days, clones were picked and checked by Western blotting.

### DNA fiber Assay

Cells were labeled with 30 µM CIdU for 30 min, washed quickly twice with PBS and exposed to 250 µM IdU for another 30 min. Cells were harvested and resuspended in PBS. Cells were then lysed with lysis buffer (200 mM TrisHCL pH7.4, 50 mM EDTA, 0.5% SDS), and DNA fibers stretched onto glass slides. The fibers were then denatured with 2.5 M HCL for 1 hr, washed with PBS and blocked with 2%BSA in PBST for 30 min. The fibers were stained with anti-BrdU antibodies recognizing CIdU and IdU. Slides were imaged at 40× on a Leica microscope.

### siRNA Transfection

C17orf53 esiRNA and control siRNA were products of Sigma. 10nM of each siRNA was transfected to cells with Lipfectamin RNAiMAX Reagent (13778, Invitrogen) according to the manufacturer’s instructions. Culture medium was changed 12 hr after the transfection.

### RNA extraction, reverse transcription and real-time PCR

RNA samples were extracted with TRIZOL reagent (Invitrogen). Reverse transcription assay was performed by using the iScript cDNA Synthesis Kit (BioRad, Hercules, CA, USA) according to the manufacturer’s instructions. Real-time PCR was performed by using Power SYBR Green PCR master mix (Applied Biosystems, Foster City, CA, USA). For quantification of gene expression, the 2 − ΔΔCt method was used. Actin expression was used for normalization.

### Immunoprecipitation (IP) and Mass Spectrometry

Expression plasmids were transfected to HEK293 suspension cells with polyethyleneimine. Cells were harvested 64 h after transfection and pellets were directly lysed with NTEN buffer (20 mM Tris-HCl [pH 7.5], 150 mM NaCl, 10% glycerol, 0.5% NP40, 10 mM NaF, 1 mM phenylmethylsulfonyl fluoride (PMSF), 1 μg ml−1 leupeptin, 1 μg ml−1 aprotinin). The lysates were ultra-centrifuged at 15,000 rpm for 15 min and then the supernatant was incubated with Streptavidin beads for 3–4 h at 4 °C. The beads were washed four times with NETN buffer and washed with NETN with 2mg/ml biotin. The elution was then incubated with S-protein beads for 2hrs. Subsequently, the eluted complexes were analyzed by sodium dodecyl sulfate– polyacrylamide gel electrophoresis (SDS-PAGE) and mass spectrometry.

### Western blotting

Cells were lysed in NET-N buffer (20 mM Tris [pH 7.6], 1 mM ethylenediaminetetraacetic acid, 1% NP40, 150 mM NaCl) supplemented with protease inhibitor cocktail tablets (Roche).

### Gel-shift Assay

The DNA substrates were made by annealing oligos 60nt-f and 60nt-r, 30nt-f and 60nt-r, respectively. The 5’ ends of the oligo 60nt-f and 30nt-f were labeled with 32P using T4 polynucleotide kinase before annealing. In all, 5 nM 32P -labeled DNA subtracts and the indicated amount of proteins were incubated at 25 °C in 10 μl reaction buffer (20 mM HEPES at pH 7.5, 5 mM MgCl2, 100 mM KCl, 1 mM DTT, 0.05% Triton X-100, 100 μg ml−1 BSA and 5% glycerol) for 15 min. Reaction mixture was loaded and resolved on a 5% Tris-borate-EDTA (TBE) gel.

### sgRNA screening

The TKOv3 library was a gift from Traver Hart lab, MD Anderson Cancer Center. DDR library was made by our lab (unpublished). The CRISPR screen was as describe before. In brief, 120 million 293A or C17orf53KO 293A cells were infected with the TKOv3 library lentiviruses or DDR library lentivirus at a low MOI (< 0.3). Twenty-four hours after infection, the medium was replaced with fresh medium containing puromycin (2 μg/mL). After selection, cells were split into 2 replicates containing #x223C;20 million cells each, passaged every 3 days, and maintained at 200-fold coverage. At day 0 and every 3 days from day 6 to day 21, 25 million cells (> 300-fold coverage) were collected for genomic DNA extraction. Genomic DNA was extracted from cell pellets using the QIAamp Blood Maxi Kit (Qiagen), precipitated using ethanol and sodium chloride, and resuspended in Buffer EB (10 mM Tris-HCl, pH 7.5). gRNA inserts were amplified via PCR using primers harboring Illumina TruSeq adapters with i5 and i7 barcodes as previously reported 38, and the resulting libraries were sequenced on an Illumina HiSeq 2500 system. The sequencing results were used for further analysis. MAGecK and Drug-Z analysis were used to calculate the difference between the DMSO- and MMC-treated groups.

### HR-GFP and NHEJ-GFP reporter Assay

U2OS cells stably expressing HR reporter DR-GFP and NHEJ reporter were gifts from Albert C. Koong lab in MD Anderson Cancer Center. 1 × 10^6^ U2OS DR-GFP or NHEJ-GFP cells were transfected with 50nmol control siRNA or C17orf53 siRNA with RNAimax. 24hr later, 2 μg of pCBASce plasmid (an I-SceI expression vector) were transfected into the cells with Lipo2000. Cells were cultured for another 48hr and subjected to flow cytometry analysis to determine percentages of GFP-positive cells, which result from HR repair or NHEJ repair induced by DNA DSBs. Means were obtained from two independent experiments.

### Luciferase based ICL repair Assay

0.2 mg/ml pCMV-Luc DNA was incubated with 5uM of cis-DDP in TE buffer (10 mM Tris-HCl pH 8.0, 1 mM EDTA) for 3h at 37°C in the dark. The reaction was stopped by NaCl (added to 0.5 M). The coss-linked Plasmid DNA (pCMV-Luc XL) was ethanol precipitated, washed in 70% ethanol, dried and redissolved in TE-buffer. 293A WT cells and *C17orf53*KO cells were transfected pCMV-Luc XL or Control (1ng of luciferease reporter plasmid/6 well plate, use 50ng of Renilla as internal control). 48hr later, cells were lysed by passive lysis buffer (Promega). Luciferase assays were performed using a dual-luciferase assay kit (Promega), quantified with monolight 3010 luminometer (BD Biosciences), and normalized to the internal Renilla luciferase control.

### Statistics

Statistical analyses were performed using GraphPad Prism software version 7.0. All of the statistical methods used are described in the main text. Each experiment was repeated twice or more, unless otherwise noted. Differences between groups were analyzed by the Student t-test. A P-value <0.05 was considered statistically significant.

## Data availability

All relevant data not presented in the main figures or Supplementary Data are available from the authors.

## Supplementary Figure Legends

**Supplementary Figure 1.**
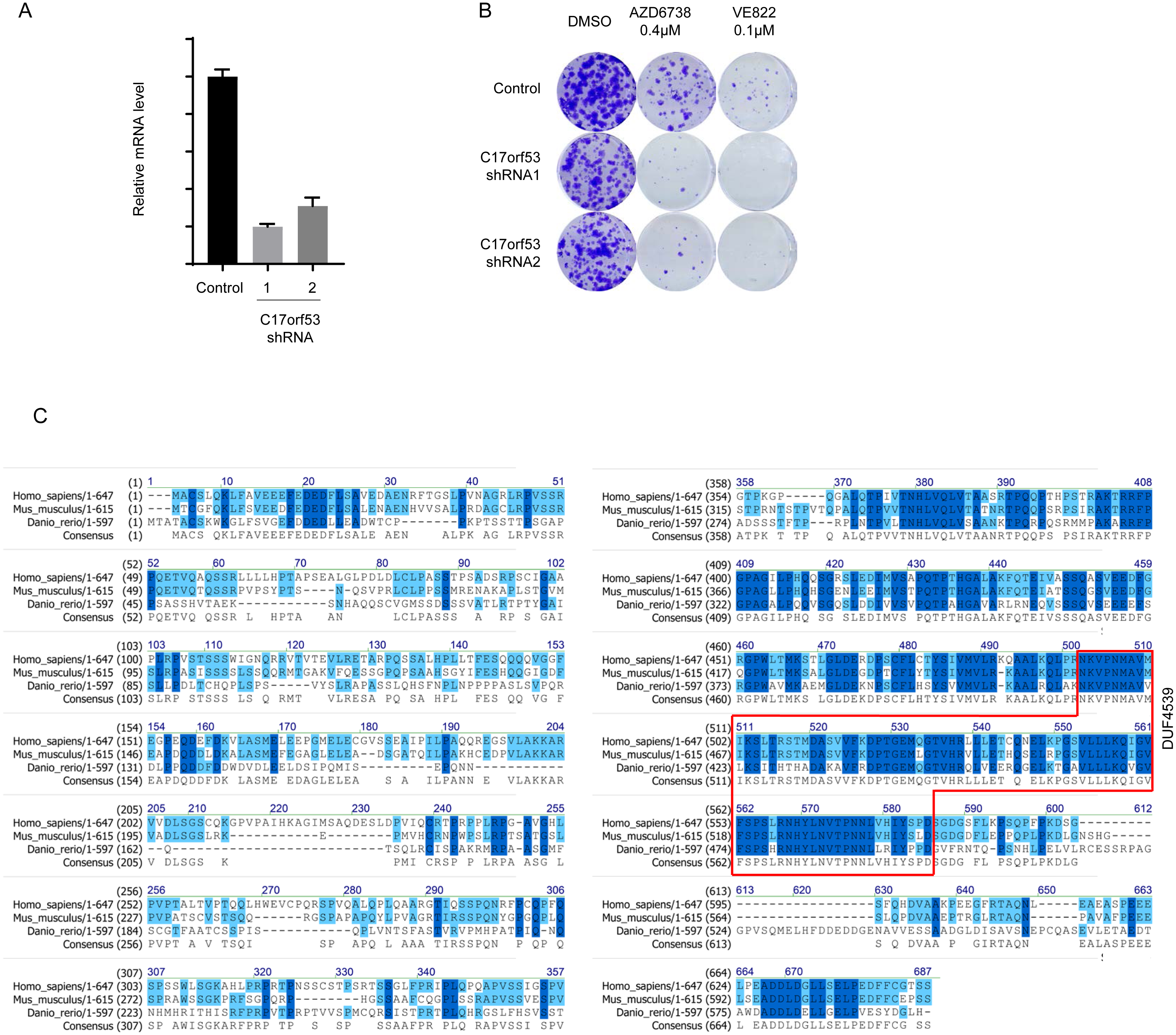
Validation of *C17orf53* as a determinant of ATRi sensitivity using shRNAs. **(A)** Realtime-PCR validation of knock-down efficiency of C17orf53 shRNAs. Mean±SD, n=2. **(B)** Representative pictures of clonogenic survival assay of cells treated with ATR inhibitors AZD6738 or VE-822. (C) Sequence alignment of C17orf53 proteins from Human, Mouse and Zebrafish.

**Supplementary Figure 2.**
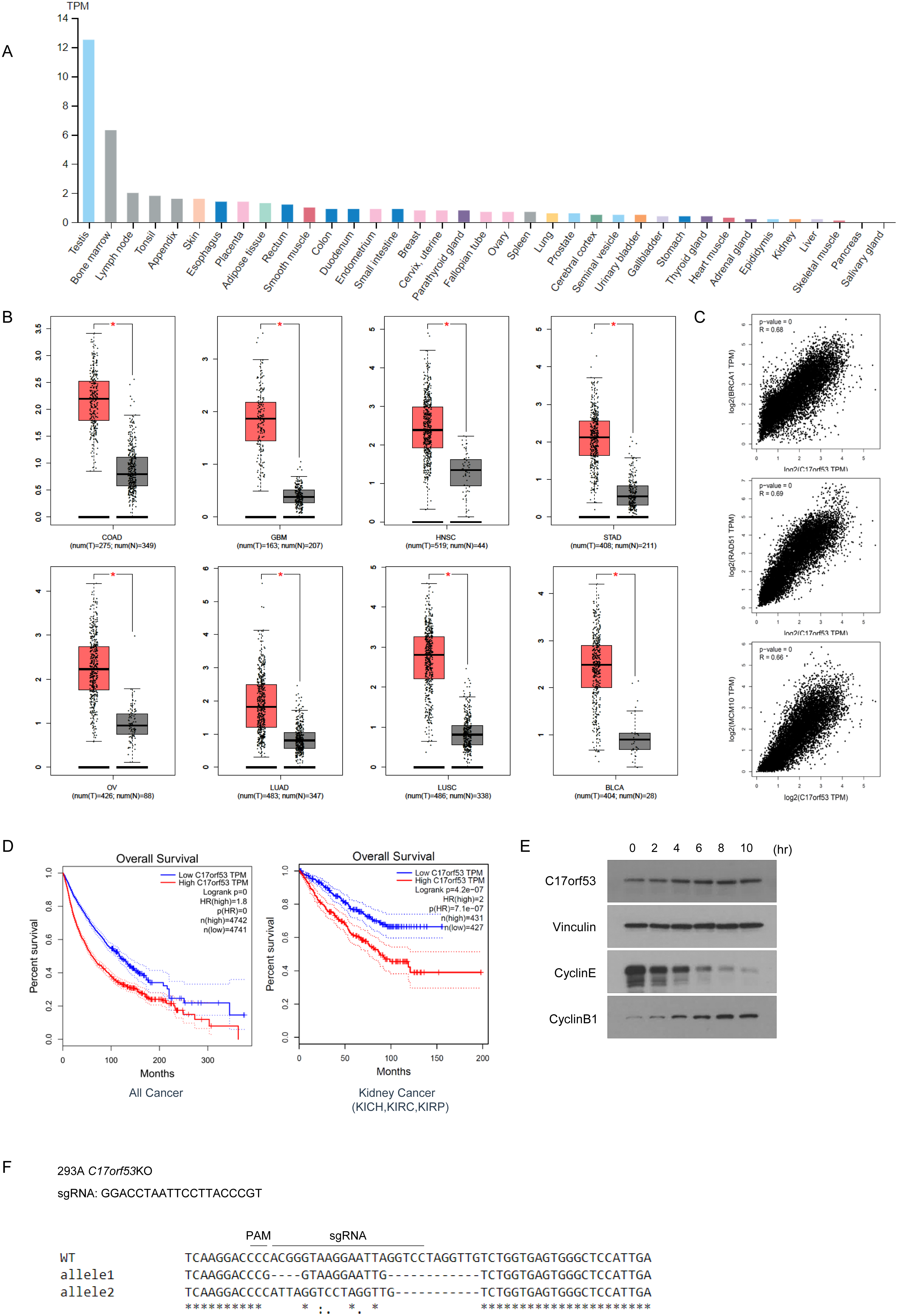
**(A)** Bar chart showing C17orf53 gene expression level across different tissues. C17orf53 expression was markedly higher in bone marrow and testis than that in other tissues. Data are from the Protein Atlas database (https://www.proteinatlas.org/). **(B)** Boxplots showing C17orf53 gene expression level between multiple types of tumors (TCGA data) and relevant normal tissues (GTEx data). C17orf53 expression was significantly higher in most of the tumors. Data were analyzed with GEPIA (http://gepia.cancer-pku.cn/). **(C)** C17orf53 expression correlates with BRCA1, RAD51 and MCM10 in tumors collected in TCGA. Data were analyzed with GEPIA (http://gepia.cancer-pku.cn/). **(D)** Overall survival for patients with tumors (more than 9000,TCGA) or pan-kidey cancers (KICH: Kidney Chromophobe; KIRC:Kidney renal clear cell carcinoma; KIRP:Kidney renal papillary cell carcinoma). Statistical significance was assessed by the log-rank test. Data were analyzed with GEPIA (http://gepia.cancer-pku.cn/). **(E)** 293A cells were synchronized by TdR treatment and released. Cells were collected at different time point. The total cell lysates were immunoblotted with the indicated antibodies. **(F)** sgRNA used to generate 293A-*C17orf53* KO cells and sequences of 293A-*C17orf53* KO cells.

**Supplementary Figure 3.**
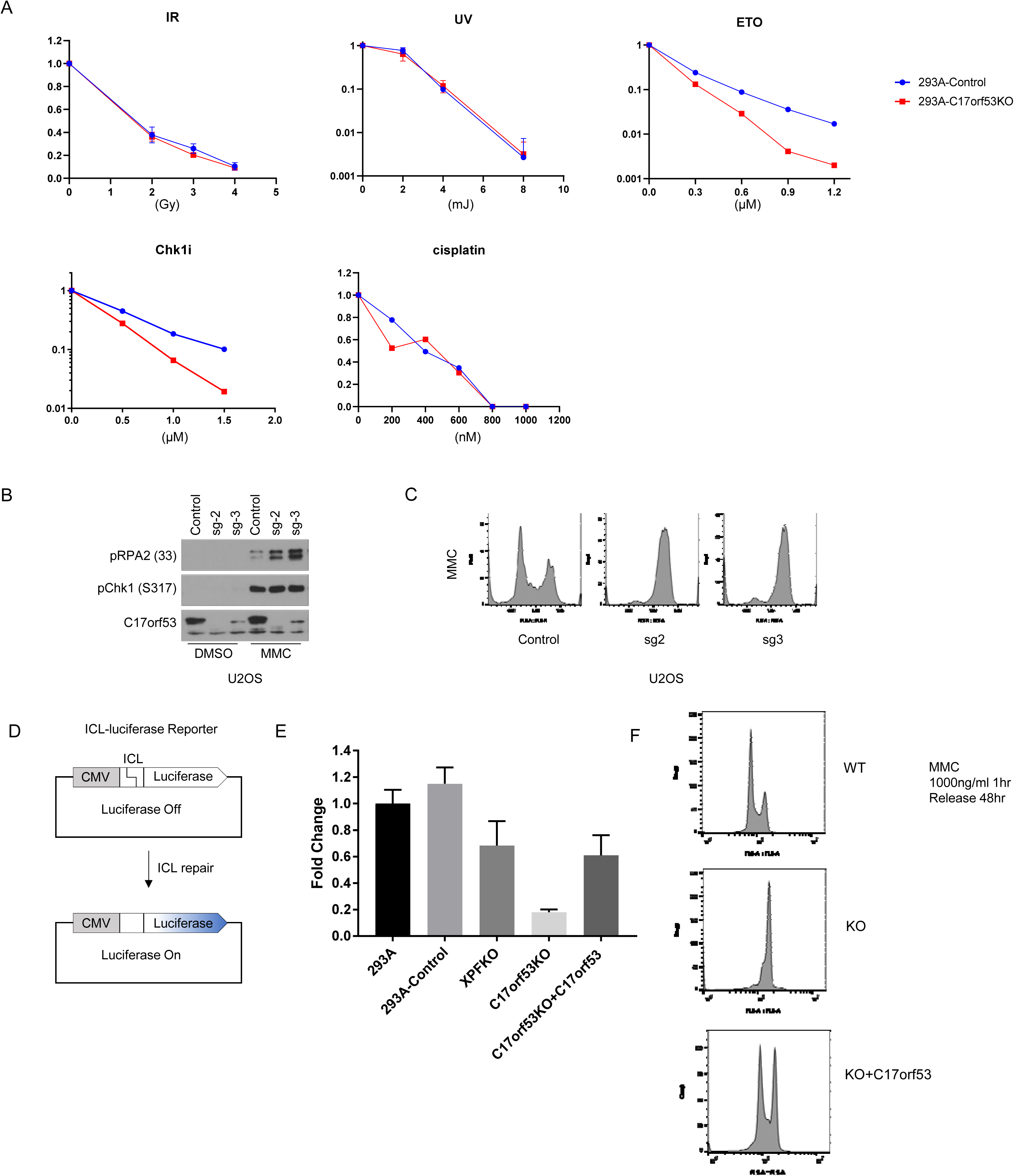
C17orf53 deficient cells show defects in ICL repair. **(A)** Clonogenic survival analysis of wild-type and C17orf53KO 293A cells exposed to indicated genotoxins. **(B)** U2OS cells infected without (control) or with different sgRNAs targeting C17orf53 were treated with DMSO or MMC (500 ng/ml, 8 hrs). Total cell lysates were immunoblotted with the indicated antibodies. **(C)** The cells in B) were treated with DMSO or MMC (10 ng/ml) for 48hrs before ethanol fixation, propidium iodide staining, and fluorescence-activated cell sorting (FACS) analysis. **(D)** Schematic scheme of luciferase-based ICL assay. **(E)** Quantification of the repair of site-specific ICL in wild-type, XPFKO, C17orf53KO and C17orf53KO+C17orf53 293A cells. **(F)** The indicated cells were treated with MMC (1000 ng/ml, 1 hr) and release for 48 hrs before ethanol fixation, propidium iodide staining, and fluorescence-activated cell sorting (FACS) analysis.

**Supplementary Figure 4.**
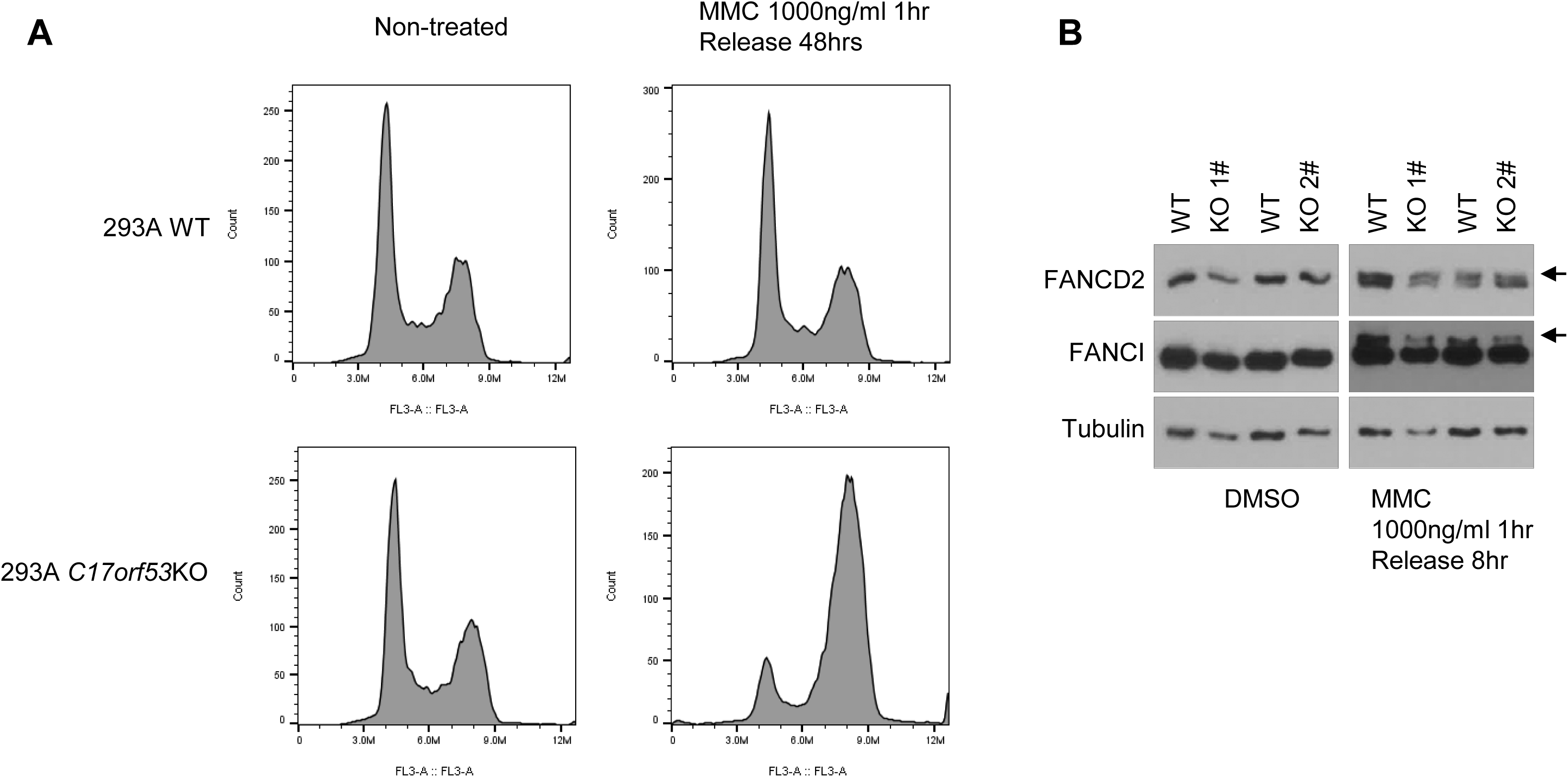
**(A)** Wild-type cells and *C17orf53*KO cells were treated with DMSO or MMC (1000 ng/ml, 1 hr) and cultured for another 48hrsbefore ethanol fixation, propidium iodide staining, and fluorescence-activated cell sorting (FACS) analysis. **(B)** MMC-induced FANCD2 and FANCI mono-ubiquitination wild-type and C17orf53 KO 293A cells. The indicated cells were left untreated or treated with MMC (1000 ng/ml, 1hr). The 8hr post-released cells were lysed and immunoblotted with the indicated antibodies.

**Supplementary Figure 5.**
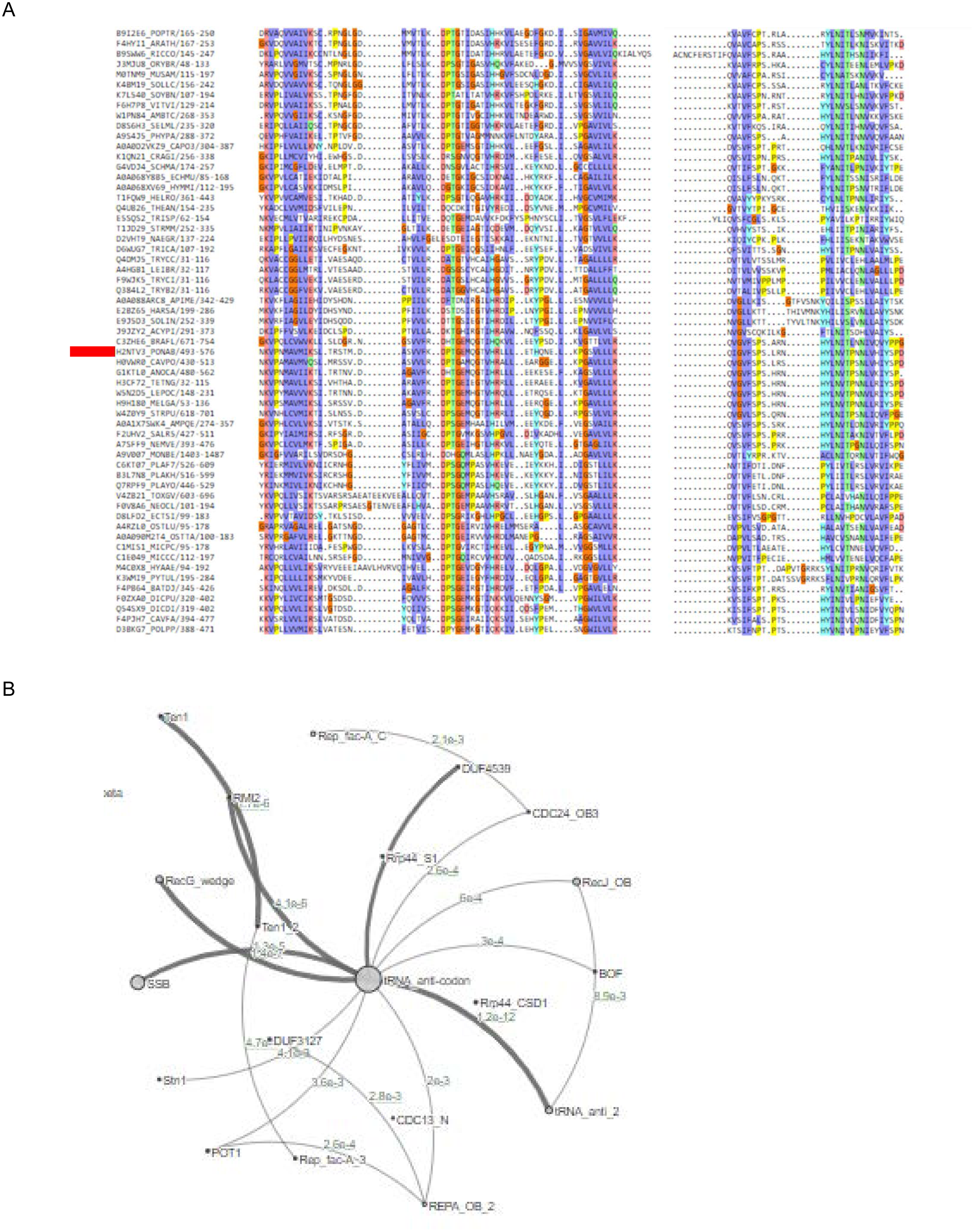
**(A)** Alignment of proteins with DUF4539 domains. (from http://pfam.xfam.org/). **(B)** Diagram shows the relationships between members of OB clan that DUF4539 belongs to. (from http://pfam.xfam.org/).

